## Supplementary Material for "FlhE functions as a chaperone to prevent formation of periplasmic flagella in Gram-negative bacteria"

**Supplementary Materials:**

Supplementary Figures S1 – S11
Supplementary Materials and Methods
Supplementary Tables S1 – S4
Supplementary References
Video S1

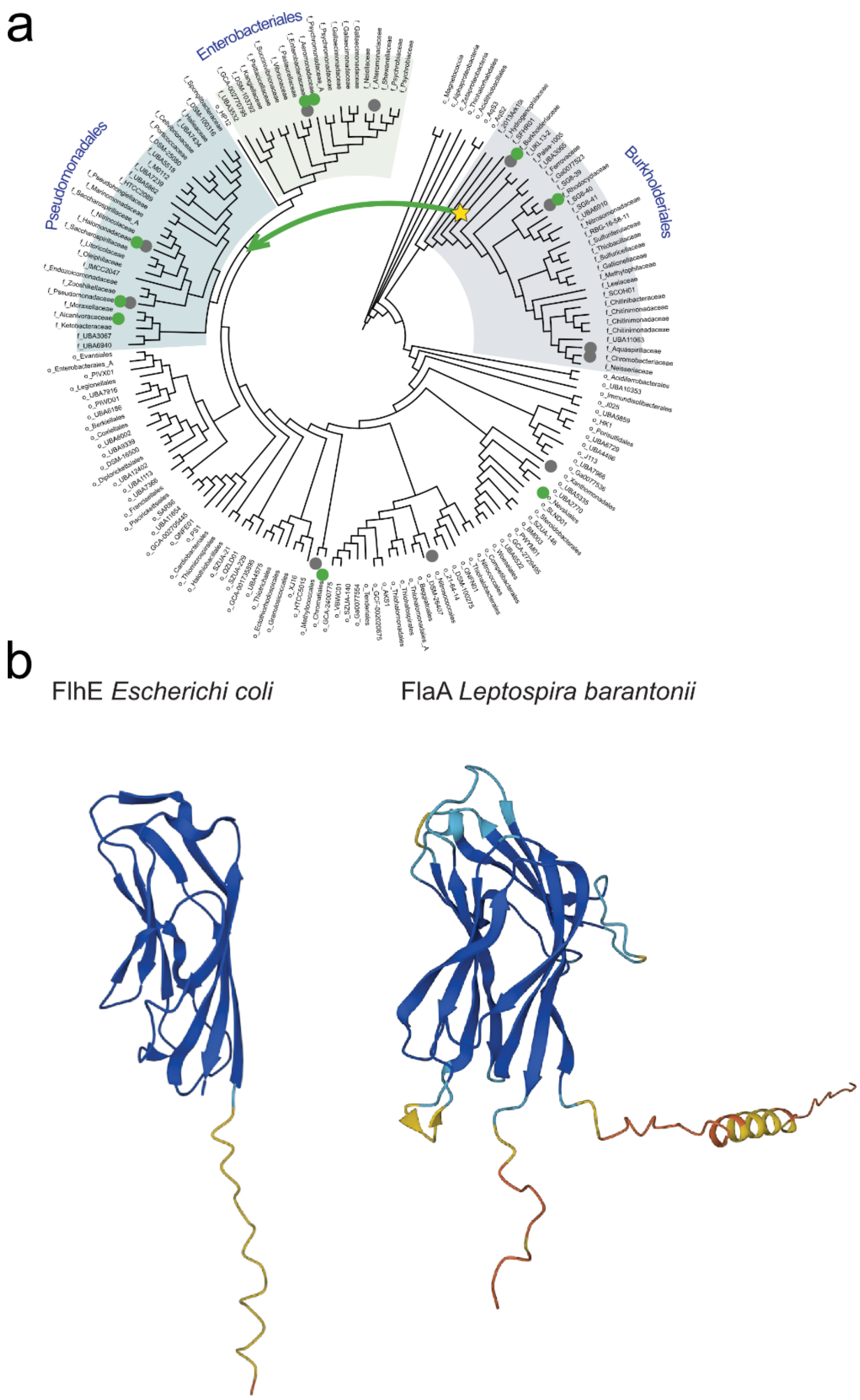


**Fig. S1:** Distribution and crystal structure of FlhE. **a** Distribution of the FlhE domain across *Gammaproteobacteria*. Green dot indicates the presence of the *flhE* gene in flagellar operons; grey dot indicates the presence of genes coding for FlhE domain containing proteins elsewhere in the genome. Star shows a plausible event of FlhE recruitment into the flagellar apparatus. Green arrow indicates horizontal gene transfer of the flagellar FlhE. **b** The crystal structure of *E. coli* FlhE (PDB: 4QXL) and the AlphaFold model of *Leptospira barantonii* FlaA (A0A5F2BH85_9LEPT) are shown. Both structures adopt the same β-sandwich fold.

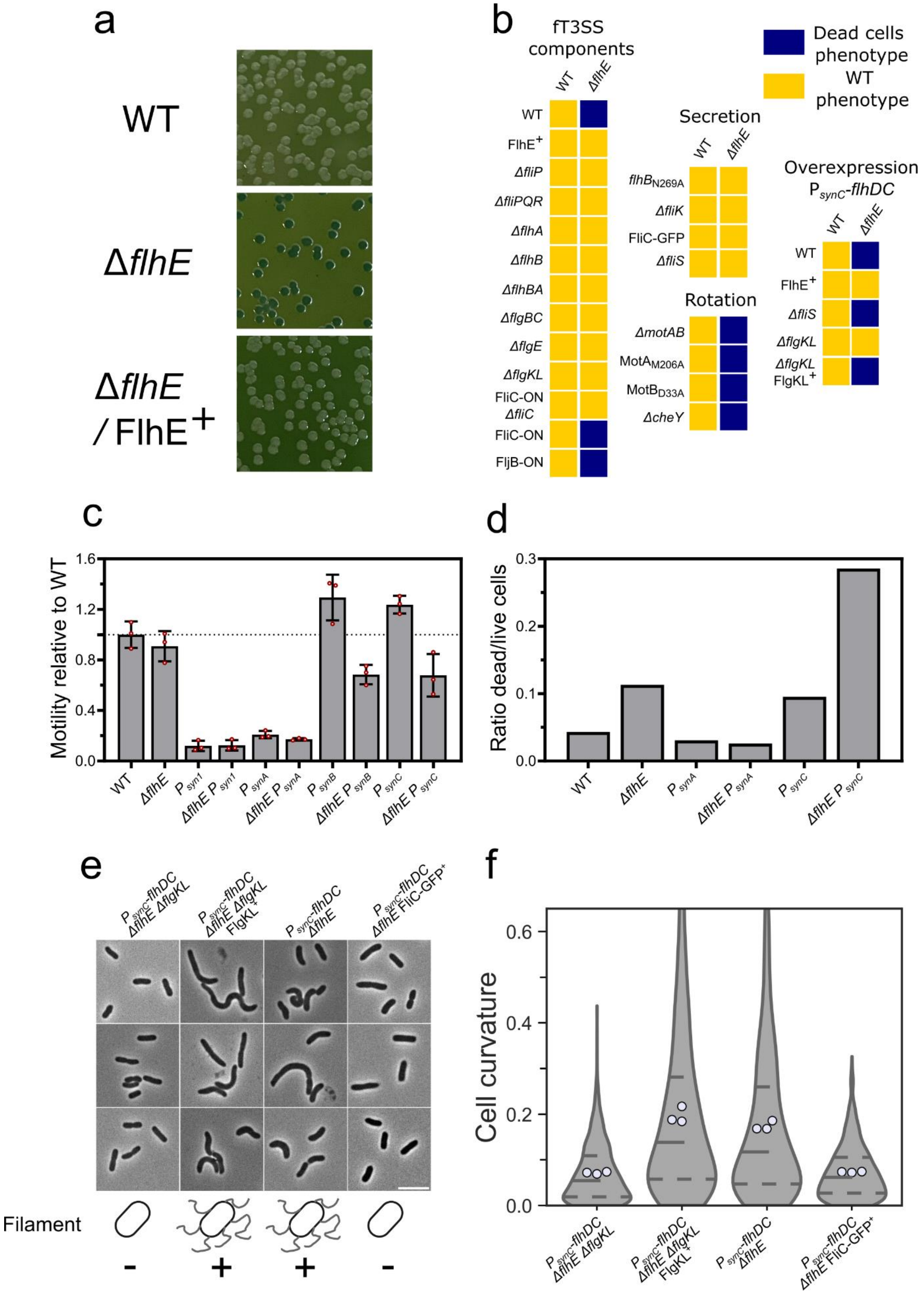
a
WT
ΔflhE
ΔflhE / FlhE+
b
fT3SS components
WT
ΔflhE
FlhE+
ΔfliP
ΔfliPQR
ΔflhA
ΔflhB
ΔflhBA
ΔflgBC
ΔflgE
ΔflgKL
FliC-ON ΔfliC
FliC-ON
FljB-ON
Secretion
flhB N269A
ΔfliK
FliC-GFP
ΔfliS
Rotation
ΔmotAB
MotA M206A
MotB D33A
ΔcheY
Dead cells phenotype
WT phenotype
Overexpression P synC-flhDC
ΔflgKL FlgKL+
c
Motility relative to WT
d
Ratio dead/live cells
e
Filament
f
Cell curvature

**Fig. S2:** Loss of FlhE causes cell death in the presence of filament formation. **a** Exemplary image of WT and Δ*flhE* mutant streaked on GP. Δ*flhE* mutant formed blue colonies compared to WT and Δ*flhE* $FlhE^{+}$ expressed constitutively *in trans* from a plasmid formed white colonies. **b** Representation of the phenotype observed on GP after deletion of genes involved in the assembly of fT3SS, secretion and flagella rotation. Yellow square represents WT phenotype and blue square cell lysis. Formation of the flagellar filament in Δ*flhE* background causes cell death. Deletion of basal body components preventing the assembly of the fT3SS restored the WT phenotype. In the presence of hook but absence of filament (FliC-ON Δ*fliC*), WT phenotype can be observed. The same phenotype is observed when blocking the secretion pore with FliC-GFP, indicating secretion itself is not responsible for the Δ*flhE* phenotype. Deletion of stator units (Δ*motAB*) or point mutants preventing the rotation ($MotA_{M206A}$ / $MotB_{D33A}$) did not restore the WT phenotype. **c** Overexpression of FlhDC causes motility decrease in Δ*flhE* background. Motility test in swimming agar were performed at 37°C. Biological triplicates were performed, and motility of the various strains was made relative to WT for each plate. Strong constitutive promoters $P_{synB}$-*flhDC* and $P_{synC}$-*flhDC* increase motility in presence of FlhE 1.5-fold relative to WT and is decreased 2-fold in Δ*flhE* background. **d** Propidium staining of WT and Δ*flhE* mutant in native promoter *flhDC*, $P_{synA}$-*flhDC* and $P_{synC}$-*flhDC* background. A steep increase in cell death can be observed in $P_{synC}$-*flhDC* Δ*flhE* background. **e** Deletion of hook-filament junction (Δ*flgKL*) and expression of FliC-GFP *in trans*, preventing the assembly of the filament and secretion of flagellins restore the WT phenotype in the $P_{synC}$-*flhDC* Δ*flhE* background. Complementation of Δ*flgKL in trans* causes the shape defect, confirming that the Δ*flgKL* deletion do not cause any polar effect on the *flg* operon. Scale bar = 5 µm. **f** Cell curvature analysis of the cells displayed in **e**. 3 independent biological replicates were performed. At least 300 cells per replicates were analysed.

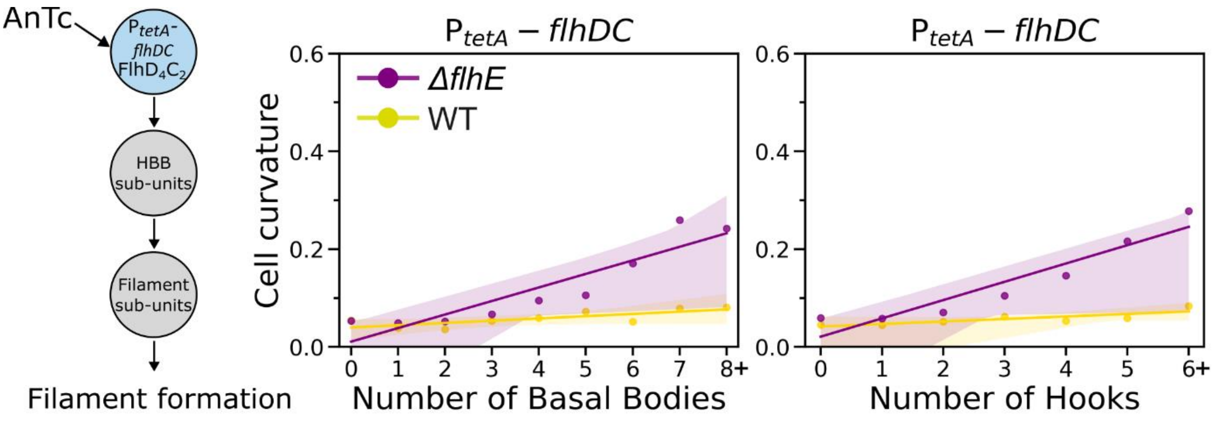


**Fig. S3:** Cell curvature increase in the Δ*flhE* mutant is correlated with an increase number of basal bodies (BB, FliG-mNeonGreen) and hooks ($FlgE_{S171C}$, Maleimide STAR RED) in an AnTc-inducible $P_{tetA}$-*flhDC* background. 3 independent biological replicates were performed. At least 250 cells per replicates were analysed using MicrobeJ. Dots represent the cell curvature mean for all cells with a given number of basal bodies or hooks. The curve represents the confidence interval, with an interval CI=99.9.

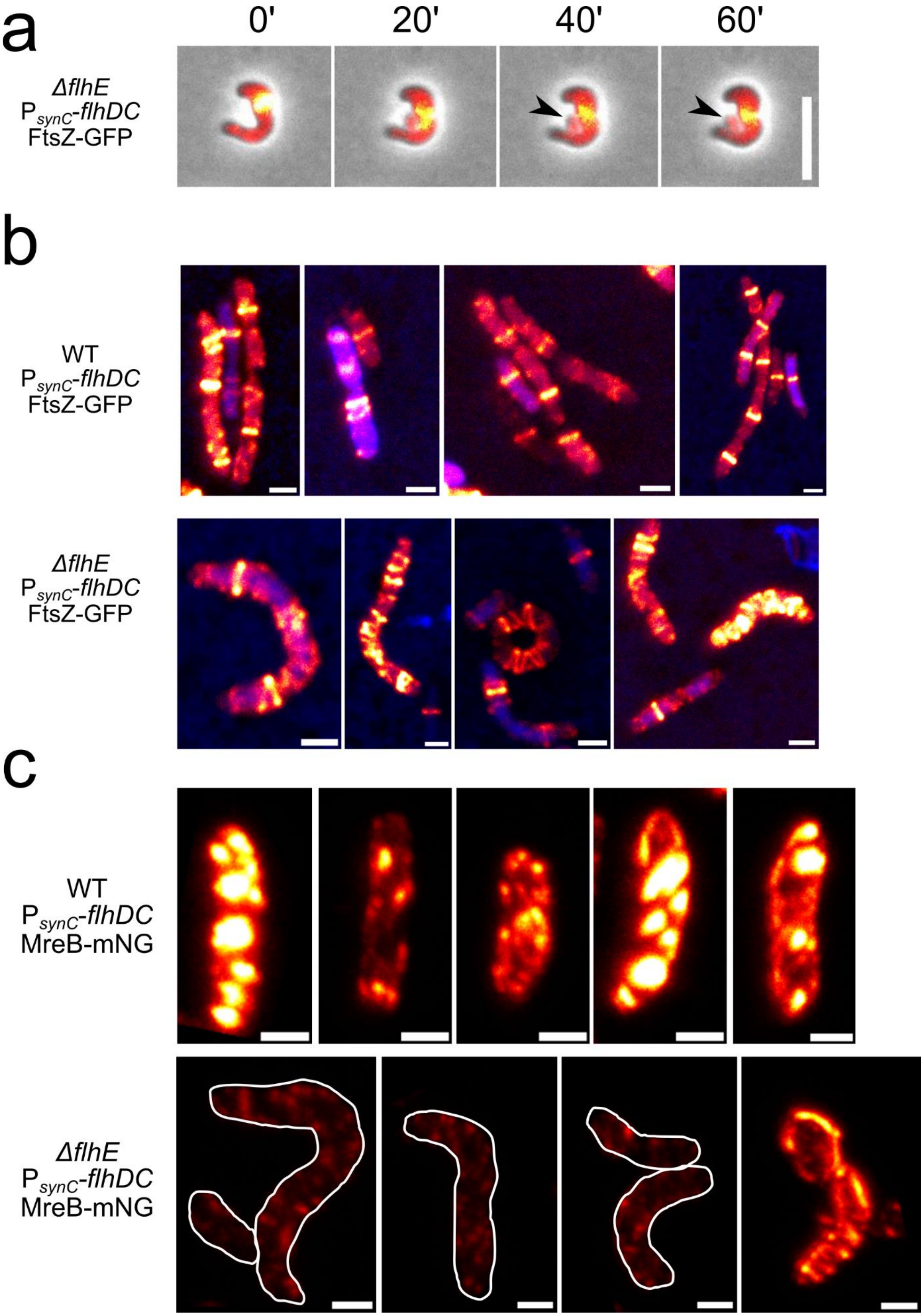


**Fig. S4**: Representative images of FtsZ-GFP and MreB-mNeonGreen. **a** Time lapse microscopy of plasmid-based expressed FtsZ-GFP in $P_{synC}$-*flhDC* Δ*flhE* mutant from overnight culture. OM rupture and release of the cytoplasm occurs at the site of the FtsZ ring position as indicated by the black arrow. Scale bar = 5 µm. **b** Exemplary image of overnight cultures of $P_{synC}$-*flhDC* and $P_{synC}$-*flhDC* Δ*flhE* mutant carrying a vector expressing FtsZ-GFP, observed by confocal microscopy. DNA staining was performed with Maleimide LIVE 560 (blue). Scale bar = 1 µm. **c** Exemplary image of overnight cultures of $P_{synC}$-*flhDC* and $P_{synC}$-*flhDC* Δ*flhE* mutant carrying a vector expressing MreB-mNeonGreen (G228-D229), observed by confocal microscopy. Scale bar = 1µm.

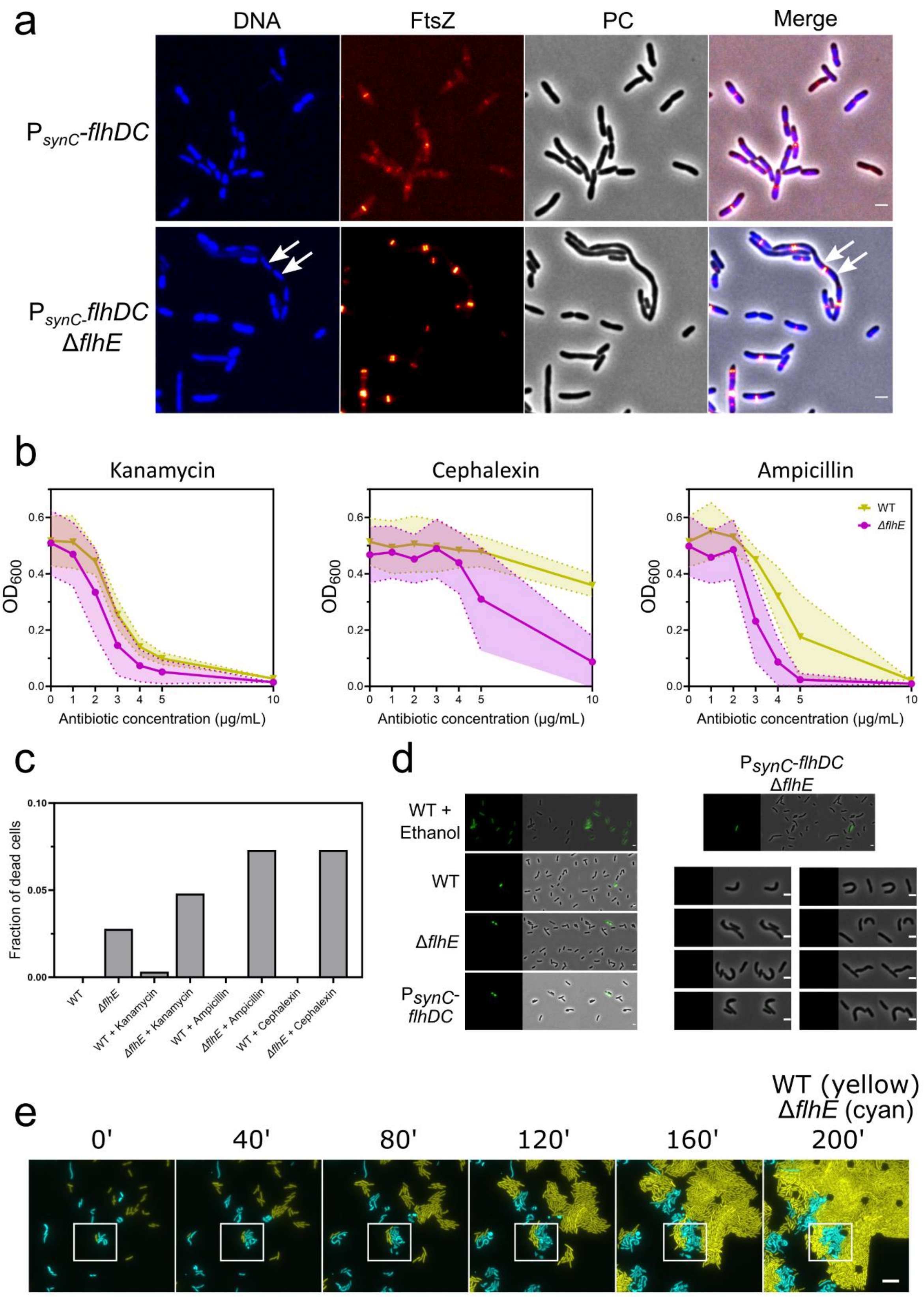


**Fig. S5:** Effect of Δ*flhE* on PG and OM. **a** DNA staining with Maleimide 560 LIVE (blue) of $P_{synC}$-*flhDC* and $P_{synC}$-*flhDC* Δ*flhE* expressing FtsZ-eGFP. Scale bar = 2 µm. **b** Killing curve of $P_{synC}$-*flhDC* and $P_{synC}$-*flhDC* Δ*flhE* in presence of different antibiotic concentration. Mid-exponential phase cultures were backdiluted to $OD_{600}$ = 0.01 in a transparent clear bottom 96-well plate. After 1 h incubation at 37°C in a plate reader, antibiotic was added at the indicated concentration. Incubation was resumed and $OD_{600}$ after 8 h incubation was plotted for each concentration. Experiment display 4 biological replicates. **c** Impact of antibiotics on cell death in $P_{synC}$-*flhDC* and $P_{synC}$-*flhDC* Δ*flhE*. Middle exponential phase cultures were treated with 10 µg/mL of antibiotics for 1.5 h at 37°C and spotted on 1% agarose pad. Cell death was visually assessed in cells losing the contrast indicative of membrane rupture and cytoplasmic contents loss. At least 300 cells per conditions were analysed. **d** DNA SYTOX GREEN staining indicate that defect in cell shape caused by Δ*flhE* is not causing OM pore formation. Left side: signal observed in a WT treated with 70% EtOH (positive control), WT, Δ*flhE* and $P_{synC}$-*flhDC* background. Right side: exemplary pictures of $P_{synC}$-*flhDC* Δ*flhE* background. Cell with aberrant morphology display no SYTOX green signal. Scale bar = 2 µm. **e** Formation of minicells in Δ*flhE* background. Microfluidic experiment (CellAsic ONIX, Merck) of $P_{synC}$-*flhDC* (yellow, mNeonGreen) and $P_{synC}$-*flhDC* Δ*flhE* (cyan, mCerulean) expressing fluorescent proteins from a constitutive promoter carried on a vector. Δ*flhE* strain display a growth defect phenotype. White square represents the field of view displayed in Fig. 2c. Scale bar = 10 µm.

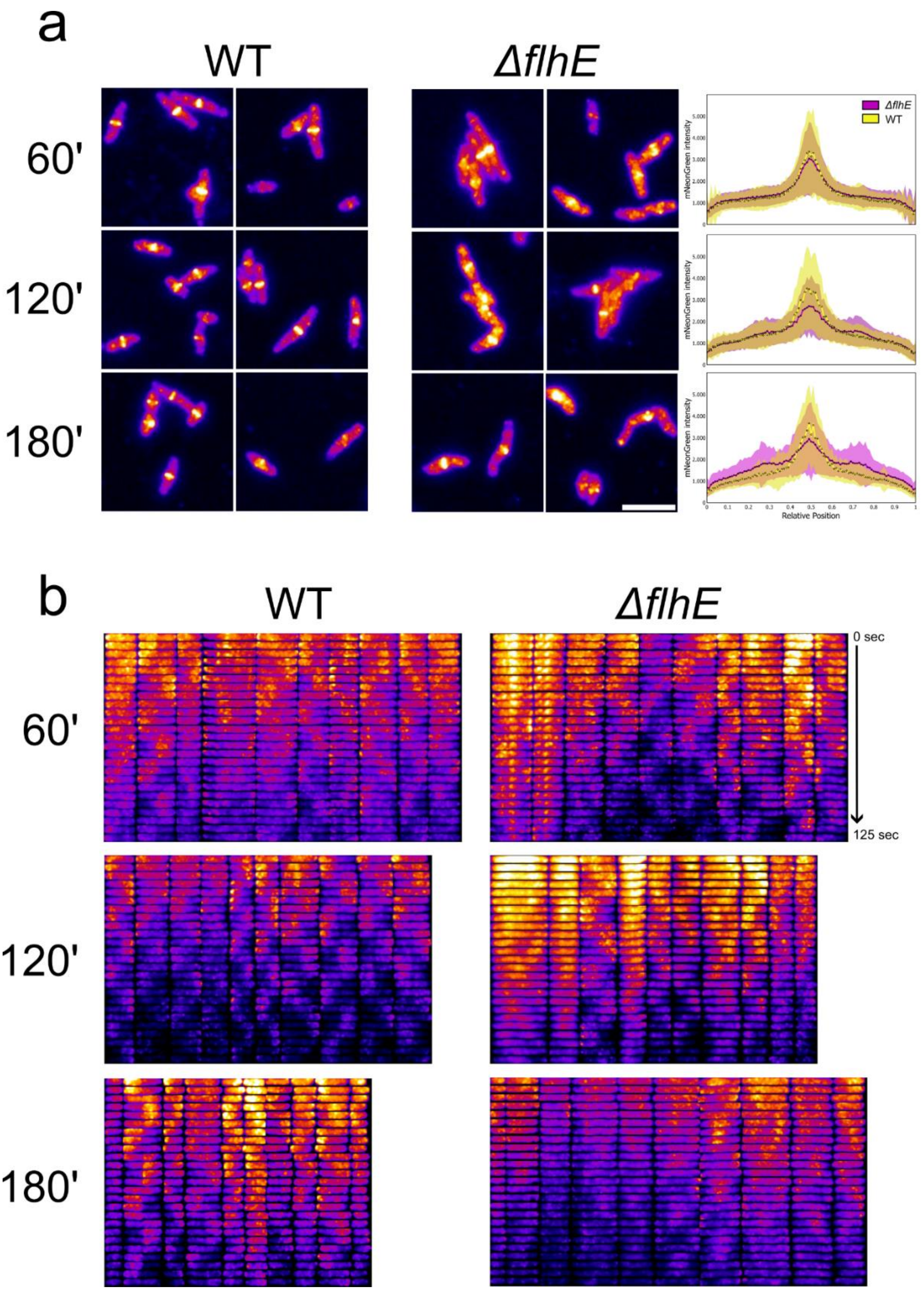


**Fig. S6:** Growth defect in Δ*flhE* Δ*flgHI* mutant perturbate ZapA localisation and MinD oscillations. **a** Time course TIRF experiment of mNeonGreen-ZapA in Δ*flgHI* and Δ*flhE* Δ*flgHI* background shows ectopic localisation of ZapA rings. Left panel: exemplary images of mNeonGreen-ZapA. Scale bar = 5 µm. Right panel: measurement of fluorescence intensity on the medial profile of at least 272 cells analysed per timepoint and strain. Overtime, the fluorescence intensity is shifting from the centre to the poles in the Δ*flhE* Δ*flgHI* background. **b** Time course TIRF experiment of mNeonGreen-MinD in Δ*flgHI* and Δ*flhE* Δ*flgHI* background. Cells were imaged 60, 120 and 180 min after AnTc induction of $P_{tetA}$-*flhDC*. A time-lapse was acquired with 5 sec intervals between each picture. Cells were aligned in a demograph using MicrobeJ. MinD oscillations is decreased in Δ*flhE* Δ*flgHI* background relative to Δ*flgHI* background.

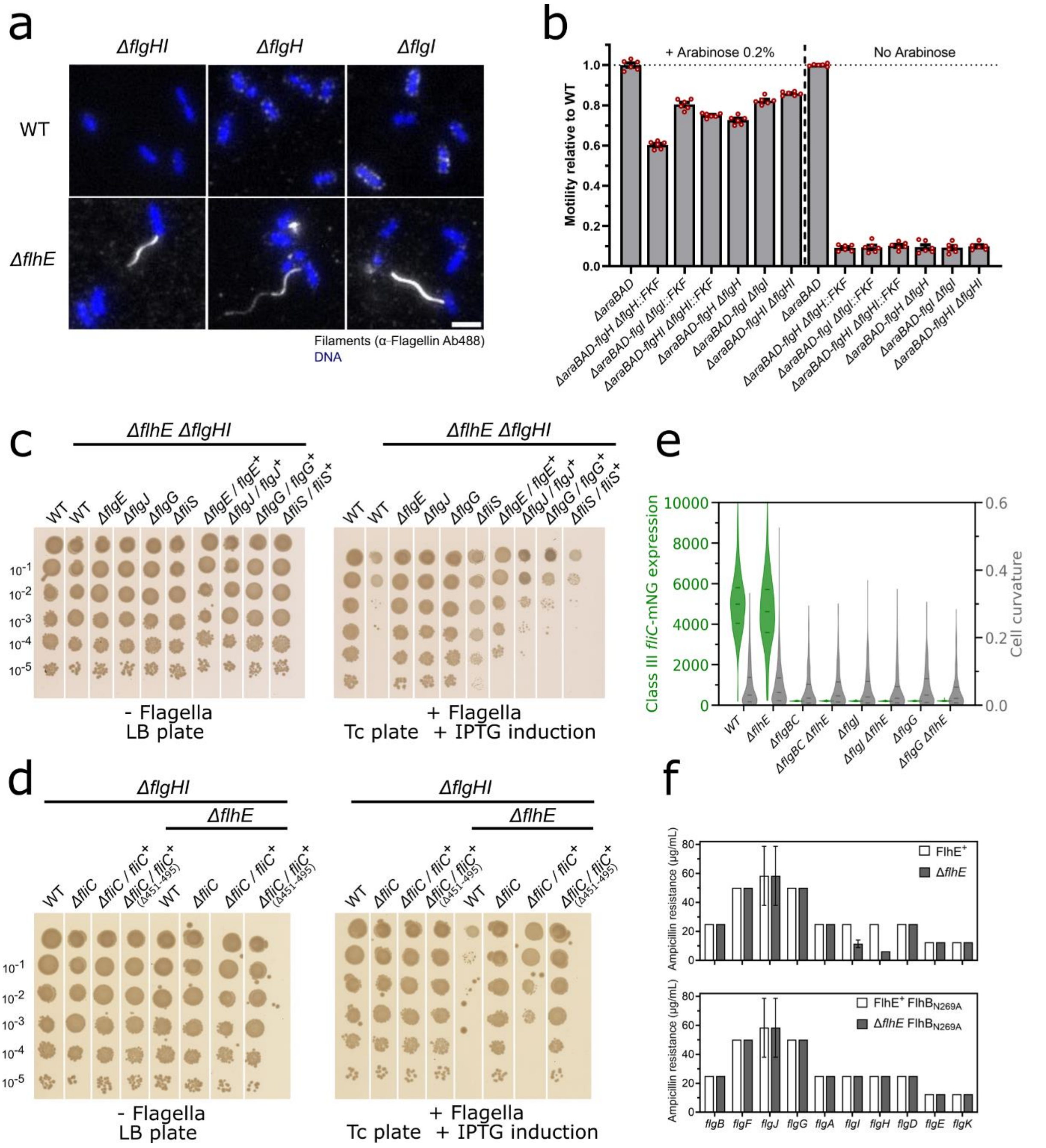

**Fig. S7:** Phenotype of Δ*flgHI* Δ*flhE* background in present of other deletion of the fT3SS structure. **a** Immunostaining of flagellar filament reveals that Δ*flhE* Δ*flgHI*/Δ*flgH*/Δ*flgI* strain can assemble filaments. Immunostaining were performed with primary anti-FliC antibodies and secondary antibodies coupled to Alexa488. DNA staining was performed with Fluoroshield with DAPI. Scale bar = 2 µm. **b** Complementation of Δ*flgHI*::FKF and Δ*flgHI* clean deletion using chromosomal *araBAD* locus showed a mild polar effect on the *flg* operon. Motility was complemented between 60 to 85%. Motility tests in swimming agar were performed at 37°C. Biological duplicates were performed with 3 technical replicates, and motility of the various strains was calculated relative to WT for each plate. **c** Spot-assay of Δ*flhE* Δ*flgHI* strains coupled with a deletion of the hook (Δ*flgE),* the rod (Δ*flgJ /* Δ*flgG),* or of the flagellin chaperone *(*Δ*fliS).* Flagellar system is only expressed in presence of tetracycline inducer (+Flagella, Tc plate). In absence of inducer (-Flagella, LB plate), no growth defect can be observed for any of the strains tested. Deletion were complemented *in trans* from an IPTG inducible vector to exclude polar effect. **d** Spot-assay of Δ*flhE* Δ*flgHI* strains coupled with a deletion of the flagellin (Δ*fliC).* Δ*fliC* was complemented *in trans* from a vector expressing the full-length FliC or a truncated C-terminal version of FliC (Δaa451-495*)* unable to assemble flagellar filaments. No growth defect could be observed in absence of full-length FliC. **e** Expression of class III reporter *fliC-mneongreen* transcriptional fusion (green violin plot) and cell curvature (grey violin plot) in WT and Δ*flhE* background (*flgHI*$^{+}$)*,* combined with a deletion of rod component *(*Δ*flgBC,* Δ*flgJ,* Δ*flgG)* 180 min after induction of flagellar system with AnTc. No class III expression can be observed in Δ*flhE* in absence of the rod components*.* Cell curvature is restored to WT in absence of the rod. At least 300 cells per strain were analysed. **f** Secretion of FliK monitored using a FliK-bla reporter in Δ*flhE* strain associated with a deletion of the rod or the PL-rings. Secretion of FliK in the periplasm is detected by ampicillin resistance from the Bla fusion. No differences in FliK secretion can be observed between WT and Δ*flhE* strain for any of the deletion, except deletion of the P/L-rings (Δ*flgH /* Δ*flgI).* Introduction of a $FlhB_{N269A}$ mutation, unable to undergo the substrate specificity switch, restore the WT levels of FliK secretion (N = 6).

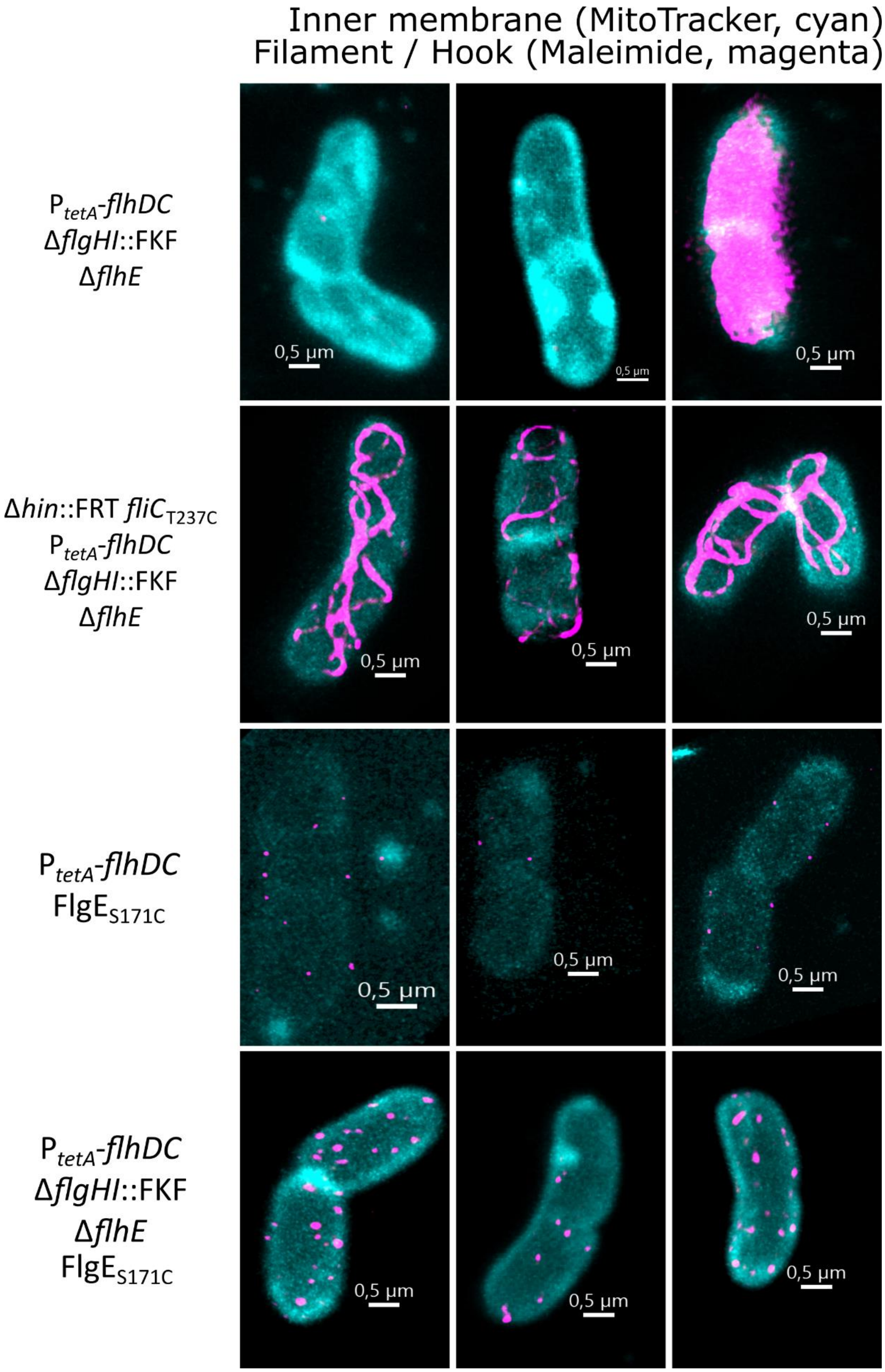


**Fig. S8:** Representative 3D images of maleimide staining of the periplasmic flagellar filament and hooks. Cells were stained with maleimide STAR RED (magenta) and MitoTracker Green (cyan) and imaged using STED super-resolution microscopy, with Z-stack intervals of 80 nm. Scale bar = 0.5 µm.

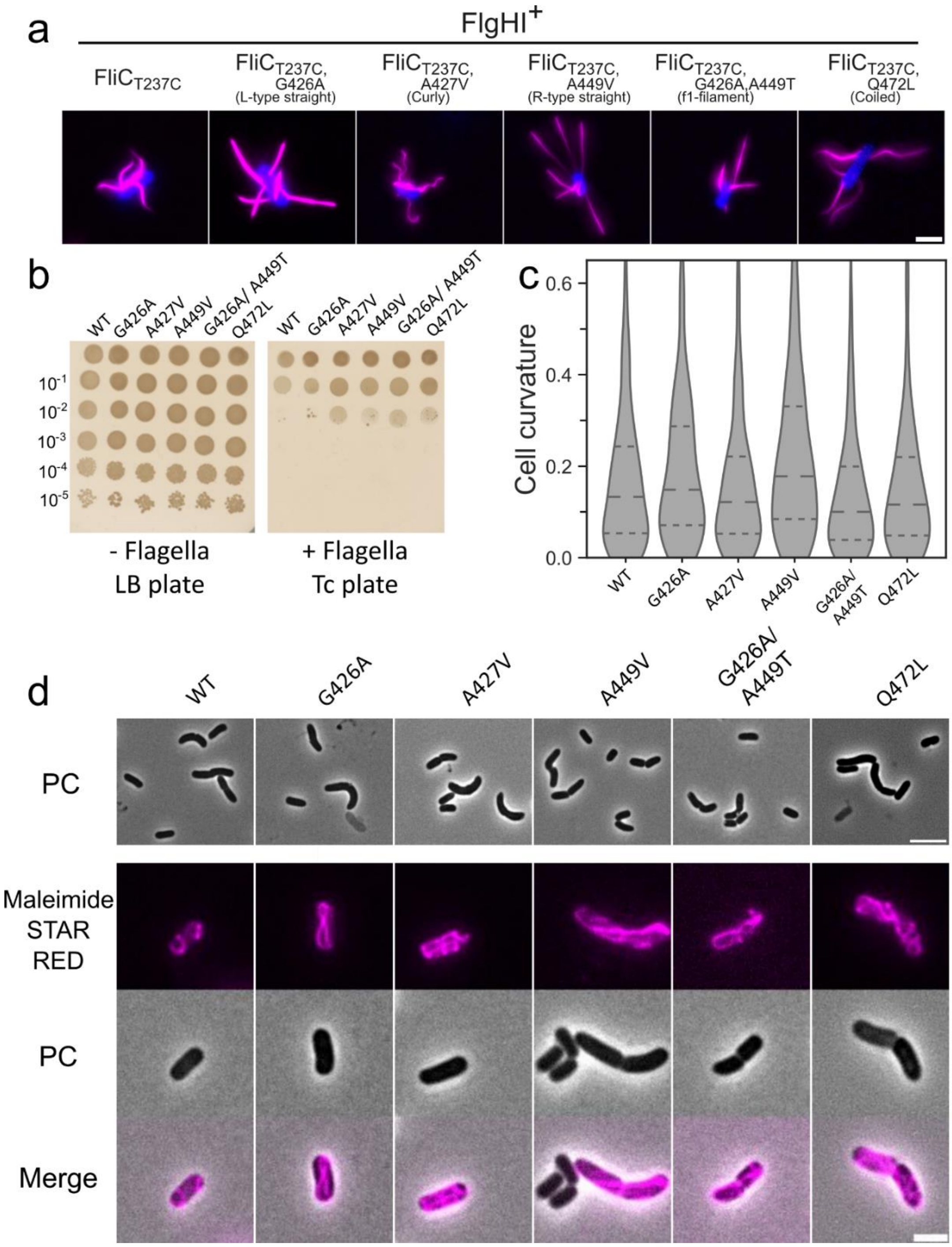


**Fig. S9:** Morphology of the flagellar filament does not impact the cell curvature. **a** Maleimide staining of $FliC_{T237C}$ with secondary point mutations changing the morphology of the filament from supercoiled to straight / curly / coiled / f1. Staining was performed in $FlgHI^+$ background, where filaments are assembled extracellularly. Scale bar = 2 µm. **b** Spot-assay of Δ*flhE* Δ*flgHI* strains coupled with the FliC point mutants described in **a**. The growth defect observed is similar between all FliC point mutants. **c** Cell curvature analysis of the strains described in **b** observed by phase contrast microscopy. At least 400 cells per strain were analysed. **d** Phase contrast and maleimide staining (magenta) of $FliC_{T237C}$ variants in Δ*flhE* Δ*flgHI background* observed by epifluorescence microscopy. Cells with curved shape can be observed independent of the FliC variant. Scale bar = 2 µm.

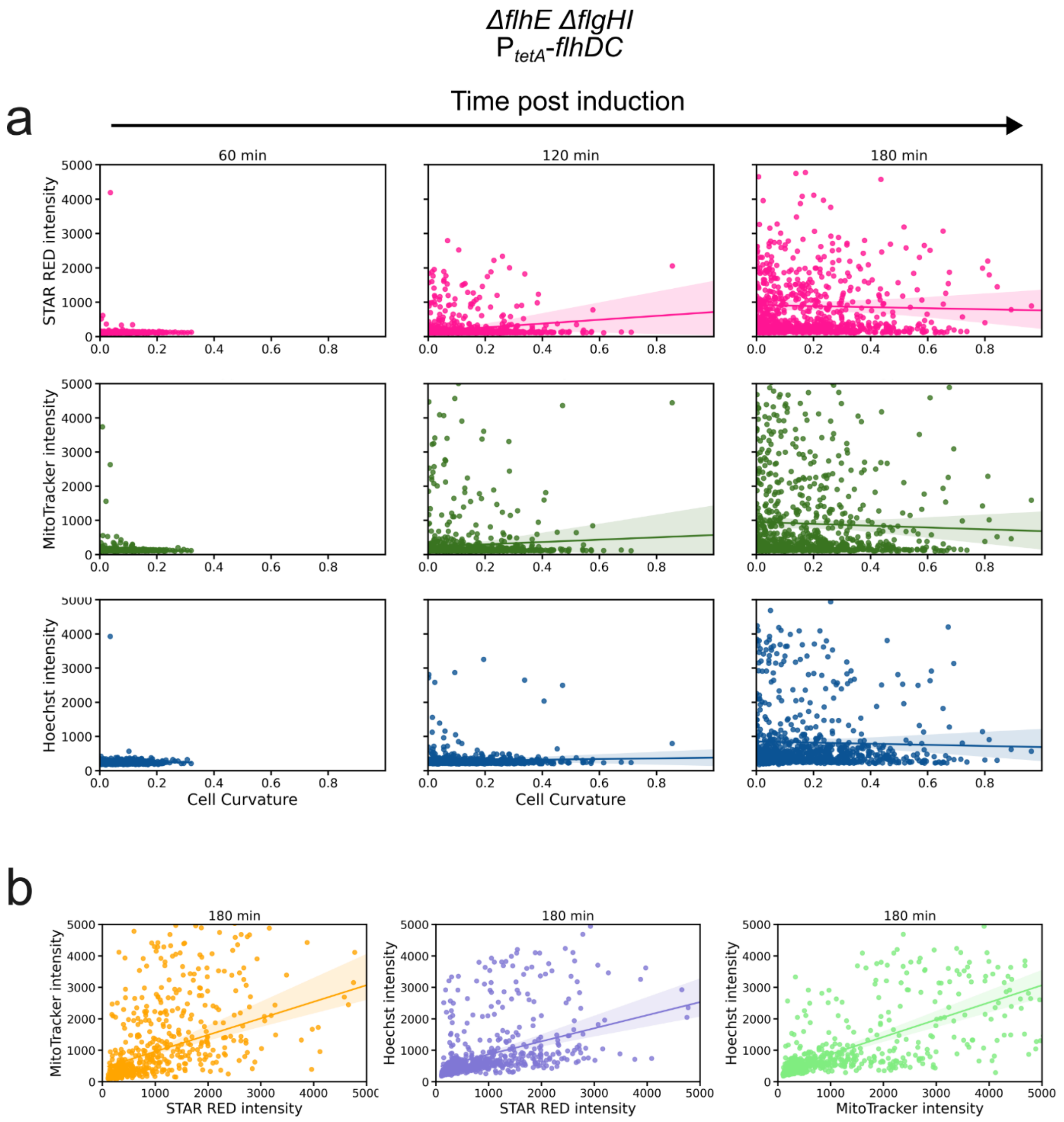


**Fig. S10:** Correlation between cell curvature and membrane staining. **a** Upper panel: STAR RED maleimide intensity plotted against cell curvature; middle panel: MitoTracker Green intensity plotted against cell curvature; lower panel: Hoechst intensity (DNA staining) plotted against cell curvature. For each staining, the increase in intensity is not directly correlated to the increase in cell curvature. **b** Dye staining relative plot. Left: STAR RED intensity plotted against MitoTracker Green; middle: STAR RED intensity plotted against Hoechst intensity; right: MitoTracker intensity plotted against Hoechst intensity. For each plot, a correlation can be observed as intensity of the different dyes increase. Biological triplicates were performed, and at least 283 cells were analysed per timepoint.

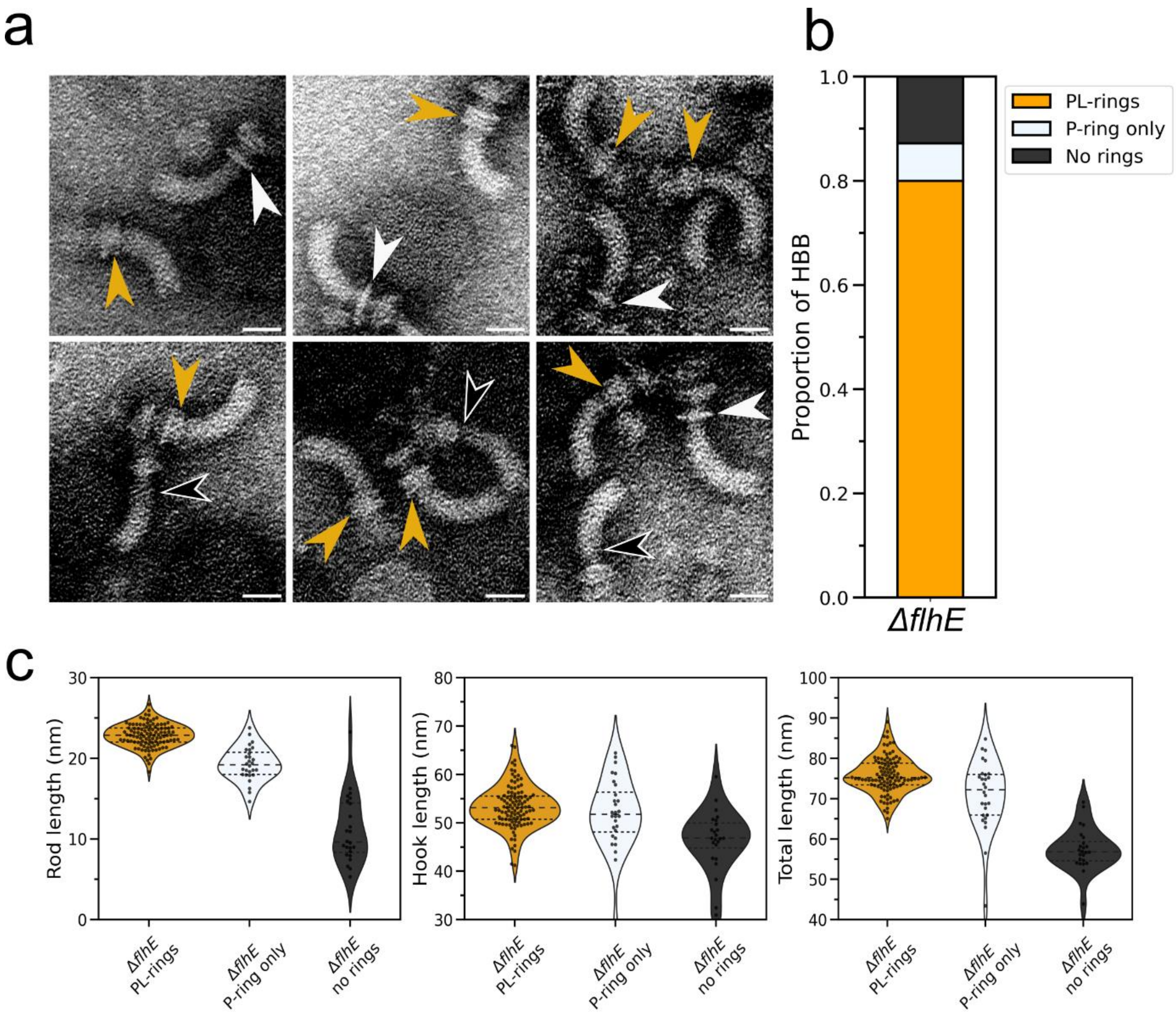


**Fig. S11:** Δ*flhE* assemble rods lacking P and L-rings. **a** Exemplary pictures of HBB purified from Δ*flhE,* observed by TEM. Most of the HBB include in their structure both PL-rings (orange arrow), while some were either missing the L-ring (P-ring only, white arrows) or both rings (no rings, black arrow). Scale bar = 25 nm. **b** Proportion of HBB with both rings, P-ring only or no rings. A total of 305 HBB were manually counted for presence of the rings. **c** Rod, hook and total (rod + hook) length for HBB from Δ*flhE* HBB purification in presence or absence of the rings. Absence of both rings is resulting in a drastic decrease of the rod length.

### Supplementary Material and Methods

#### Construction vector MreB-mNeonGreen / mNeonGreen-MinD

Plasmids pEM12582 (MreB-mNeonGreen) and pEM12581 (MinD-mNeonGreen) were constructed as follow. Overlapping PCR is a variant of standard PCR, consisting in assembling several fragments amplified by PCR in one reaction. Sequences of interests were amplified separately with Q5 High-fidelity polymerase, with oligonucleotides with flanking regions homologous to the forward / reverse fragment (primers 5401 to 5406 for *mreB*, 5409 to 5412 for *minD* in Table S5). Fragments were gel-purified using the NucleoSpin® Gel and PCR Clean-up Kit (Macherey-Nagel). PCR products were mixed together in a ratio 1:1 with Q5 PCR mix in absence of primers and incubated in the PCR thermocycler. Initial 10 cycles enable amplifications of the final construct using the PCR products and their flanking regions to polymerise the final fragment length. After 10 cycles, PCR was paused, 0.5 µM of primers amplifying from the 5'-end of first fragment to 3'-end of the last fragment were added. PCR was resumed for 25 additional cycles. Amplification was checked by electrophoresis and correct fragment was purified using the NucleoSpin® Gel and PCR Clean-up Kit (Macherey-Nagel) for subsequent Gibson Assembly. Plasmids were constructed by Gibson Assembly using Gibson Assembly Master Mix or HiFi Master Mix purchased from NEB. The vector was linearized by PCR (primers 5407/5408 for *mreB*, 5413 to 5414 for *minD* in Table S5)and gel purified after treatment with DpnI (NEB) for 2 - 4 h at 37 °C to digest methylated plasmid template DNA. The insert was amplified with specific primers containing 20 - 40 bp overhangs to the linearized vector insertion site. The Gibson Assembly reaction was performed using 0.02 - 0.05 pmol linearized vector and 0.1 - 0.2 pmol purified insert. The samples were incubated for 1 h at 50 °C before cleaning (NucleoSpin® PCR Clean-up Kit (MachereyNagel)) and transformation of 3 µL of purified plasmid into electrocompetent *S. enterica* serovar Typhimurium cells prepared freshly. Transformants were spread on selective LB agar, checked by PCR and positive PCR products were sent to sequencing after purification. MinD-mNeonGreen was then integrated in the chromosomal native locus using the λ-RED system.

### **Table S1**: Strains used in this study

| Name | Genotype | Reference |
|---|---|---|
| TH437 | LT2 | J. Roth |
| TH12483 | Δ*flhE*7404 | Lab collection |
| EM9060 | Δ*flhE7404* / pEM8913 (pKH70-P$_{rpsM}$-*flhE*) | This study |
| EM11563 | Δ*flhE*23076::FCF | This study |
| EM12322 | Δ*flhE*23076::FRT | This study |
| EM11990 | Δ*flhE*7404 Δ*fliP*23466::FRT | This study |
| EM11991 | Δ*flhE*7404 Δ*fliPQR*23467::FRT | This study |
| EM11988 | Δ*flhE*7404 Δ*flhA*23464::FRT | This study |
| EM11987 | Δ*flhE*7404 Δ*flhB*23463::FRT | This study |
| EM11989 | Δ*flhE*7404 Δ*flhBA*23465::FRT | This study |
| EM11028 | Δ*flhE*7404 Δ*flgBC*6557 | This study |
| EM8344 | Δ*flhE*7404 Δ*flgE*22964::FCF | This study |
| EM11243 | Δ*flhE*7404 Δ*hin*5717::FRT Δ*flgKL*5739::FKF | This study |
| EM12631 | Δ*flhE*7404 Δ*flgKL*7770 | This study |
| EM11557 | Δ*flhE*7404 Δ*fliC*7861::FRT Δ*hin*-5717::FCF (fliC$^{ON}$) | This study |
| EM8498 | Δ*flhE*7404 Δ*hin*-5717::FRT (fliC$^{ON}$) | This study |
| EM11556 | Δ*flhE*7404 Δ*fliC*7861::FRT Δ*hin*-5718::FCF (fljB$^{ON}$) | This study |
| EM8397 | Δ*flhE*7404 *flhB*23151$_{N269A}$ | This study |
| EM12336 | *fliK*6620(Δ aa 248-298) Δ*flhE*23076::FCF | This study |
| EM12337 | Δ*hin*-5717::FRT *flgE*6506$_{S171C}$ Δ*fliK*6137 (clean deletion) Δ*flhE*23076::FCF | This study |
| EM8493 | Δ*flhE*7404 Δ*fliS*5728::FRT | This study |
| EM12379 | Δ*hin*-5717::FRT *fliC*5936(FliC-GAGAGAGA-GFPmut2) Δ*flhE*23076::FCF | This study |
| EM12130 | Δ*flhE*23076::FCF Δ*motAB*::tetRA (Δaa R90-F63) | This study |
| EM12516 | Δ*motAB* (leaving first and last 15 bp) Δ*flhE*23076::FCF | This study |
| EM12515 | *motA*5461::*mudJ* Δ*flhE*23076::FCF | This study |
| EM12487 | *motA*$_{M206A}$ Δ*flhE*23076::FCF | This study |
| EM12489 | *motB*$_{D33A}$ Δ*flhE*23076::FCF | This study |
| EM12008 | Δ*flhE*23076::FCF Δ*cheY* 22371::FKF | This study |
| EM9659 | Δ*hin*-5717::FRT *flgE*6506$_{S171C}$ P$_{proC}$-*flhDC*23252 (Δbp -598 to AUG of *flhD)* | Lab collection |
| EM10697 | Δ*hin*-5717::FRT *flgE*6506$_{S171C}$ P$_{proC}$-*flhDC*23252 Δ*flhE*23076::FCF | This study |
| EM12325 | Δ*hin*-5717::FRT *flgE*6506$_{S171C}$ P$_{proC}$-*flhDC*23252 Δ*flhE*23076::FRT | This study |
| EM12439 | Δ*hin*-5717::FRT *flgE*6506$_{S171C}$ P$_{proC}$-*flhDC*23252 / pEM8913 (pKH70-P$_{rpsM}$-*flhE*) | This study |
| EM12440 | Δ*hin*-5717::FRT *flgE*6506$_{S171C}$ P$_{proC}$-*flhDC*23252 Δ*flhE*23076::FRT / pEM8913 (pKH70-P$_{rpsM}$-*flhE*) | This study |
| EM12765 | Δ*hin*-5717::FRT *flgE*6506$_{S171C}$ P$_{proC}$-*flhDC*23252 Δ*fliS*5728::FKF | This study |
| EM12766 | Δ*hin*-5717::FRT *flgE*6506$_{S171C}$ P$_{proC}$-*flhDC*23252 Δ*flhE*23076::FRT Δ*fliS*5728::FKF | This study |
| EM13389 | Δ*hin*-5717::FRT *flgE*6506$_{S171C}$ P$_{proC}$-*flhDC*23252 Δ*flgKL*5739::FKF | This study |
| EM12366 | Δ*hin*-5717::FRT *flgE*6506$_{S171C}$ P$_{proC}$-*flhDC*23252 Δ*flhE*23076::FCF Δ*flgKL*5739::FKF | This study |
| EM13681 | Δ*hin*-5717::FRT *flgE*6506$_{S171C}$ P$_{proC}$-*flhDC*23252 Δ*flgKL*5739::FKF Δ*araBAD*1404::*flgKL* | This study |
| EM13744 | Δ*hin*-5717::FRT *flgE*6506$_{S171C}$ P$_{proC}$-*flhDC*23252 Δ*flgKL*5739::FKF Δ*araBAD*1404::*flgKL* Δ*flhE*23076::FCF | This study |
| EM9662 | Δ*hin*-5717::FRT *flgE*6506$_{S171C}$ P$_{pro1}$-*flhDC*23252 (Δbp -598 to AUG of *flhD)* | Lab collection |
| EM9661 | Δ*hin*-5717::FRT *flgE*6506$_{S171C}$ P$_{proA}$-*flhDC*23252 (Δbp -598 to AUG of *flhD)* | Lab collection |
| EM9660 | Δ*hin*-5717::FRT *flgE*6506$_{S171C}$ P$_{proB}$-*flhDC*23252 (Δbp -598 to AUG of *flhD)* | Lab collection |
| EM10165 | Δ*hin*-5717::FRT *flgE*6506$_{S171C}$ P$_{pro1}$-*flhDC*23252 Δ*flhE*23076::FCF | This study |
| EM10164 | Δ*hin*-5717::FRT *flgE*6506$_{S171C}$ P$_{proA}$-*flhDC*23252 Δ*flhE*23076::FCF | This study |
| EM10698 | Δ*hin*-5717::FRT *flgE*6506$_{S171C}$ P$_{proB}$-*flhDC*23252 Δ*flhE*23076::FCF | This study |
| EM13743 | Δ*hin*-5717::FRT *flgE*6506$_{S171C}$ P$_{proC}$-*flhDC*23252 Δ*araBAD*1405::*fliC*23546-GAGAGAGA-GFPmut2 Δ*flhE*23076::FCF | This study |
| EM8250 | *fliG*22799 (mNeonGreen-FliG, N-terminal) *flgE*6506$_{S171C}$ | Lab collection |
| EM11018 | *fliG*22799 *flgE*6506$_{S171C}$ Δ*flhE*23076::FCF | This study |
| EM15307 | *fliG*22799 *flgE*6506$_{S171C}$ P$_{tetA}$*flhDC*5451::Tn10dTc[del-25] | This study |
| EM15308 | *fliG*22799 *flgE*6506$_{S171C}$ Δ*flhE*23076::FCF P$_{tetA}$*flhDC*5451::Tn10dTc[del-25] | This study |
| EM12501 | Δ*hin*-5717::FRT *flgE*6506$_{S171C}$ P$_{proC}$-*flhDC*23252 *fliG*22799 | This study |
| EM12844 | Δ*hin*-5717::FRT *flgE*6506$_{S171C}$ P$_{proC}$-*flhDC*23252 *fliG*22799 Δ*flhE*23076::FCF | This study |
| EM12367 | DH5alpha / pXY027 (pCA24N-*ftsZ-GFP*) | Addgene Plasmid #98915 [1] |
| EM12437 | Δ*hin*-5717::FRT *flgE*6506$_{S171C}$ P$_{proC}$-*flhDC*23252 / pXY027 (pCA24N-*ftsZ-GFP*) | This study |
| EM12438 | Δ*hin*-5717::FRT *flgE*6506$_{S171C}$ P$_{proC}$-*flhDC*23252 Δ*flhE*23076::FRT / pXY027 (pCA24N-*ftsZ-gfp*) | This study |
| EM12594 | Δ*hin*-5717::FRT *flgE*6506$_{S171C}$ P$_{proC}$-*flhDC*23252 / pEM12582 (pCA24N-*mreB*(G228)-*mneongreen-mreB*(D229)) | This study |

| | | |
|---|---|---|
| EM12597 | Δ*hin*-5717::FRT *flgE*6506$_{S171C}$ P$_{proC}$-*flhDC*23252 Δ*flhE*23076::FRT / pEM12582 (pCA24N-*mreB*(G228)-*mneongreen*-*mreB*(D229)) | This study |
| EM12362 | Δ*hin*-5717::FRT *flgE*6506$_{S171C}$ P$_{proC}$-*flhDC*23252 / pEM8731 (pKH70-P$_{rpsM}$-*mneongreen*) | This study |
| EM12550 | Δ*hin*-5717::FRT *flgE*6506$_{S171C}$ P$_{proC}$-*flhDC*23252 Δ*flhE*23076::FRT / pEM8313 (pKH70-P$_{rpsM}$-*mcerulean*) | This study |
| TH3730 | P$_{tetA}$*flhDC*5451::Tn10dTc[del-25] | Lab collection |
| EM10212 | P$_{tetA}$*flhDC*5451::Tn10dTc[del-25] Δ*flhE*23076::FCF | This study |
| TH14357 | Δ*flgBC*6557 P$_{tetA}$*flhDC*5451::Tn10dTc[del-25] | Lab collection |
| EM14564 | Δ*flgBC*6557 P$_{tetA}$*flhDC*5451::Tn10dTc[del-25] Δ*flhE*23076::FCF | This study |
| EM14079 | Δ*flgHI*958 P$_{tetA}$*flhDC*5451::Tn10dTc[del-25] | This study |
| EM14216 | Δ*flgHI*958 P$_{tetA}$*flhDC*5451::Tn10dTc[del-25] Δ*flhE*23076::FCF | This study |
| EM14242 | Δ*flgH*7662 P$_{tetA}$*flhDC*5451::Tn10dTc[del-25] | This study |
| EM14217 | Δ*flgH*7662 P$_{tetA}$*flhDC*5451::Tn10dTc[del-25] Δ*flhE*23076::FCF | This study |
| EM14243 | Δ*flgI*7663 P$_{tetA}$*flhDC*5451::Tn10dTc[del-25] | This study |
| EM14218 | Δ*flgI*7663 P$_{tetA}$*flhDC*5451::Tn10dTc[del-25] Δ*flhE*23076::FCF | This study |
| EM9357 | *fliC*23165-*mneongreen* (transcriptional fusion, insertion RBS (P$_{rpsM}$) in 3'UTR *fliC*) P$_{tetA}$*flhDC*5451::Tn10dTc[del-25] Δ*hin*-5717::FRT | This study |
| EM13604 | *fliC*23165-*mneongreen* P$_{tetA}$*flhDC*5451::Tn10dTc[del-25] Δ*hin*-5717::FRT Δ*flhE*23076::FRT | This study |
| EM14435 | *fliC*23165-*mneongreen* P$_{tetA}$*flhDC*5451::Tn10dTc[del-25] Δ*hin*-5717::FRT Δ*flgHI*23618::FKF | This study |
| EM14436 | *fliC*23165-*mneongreen* P$_{tetA}$*flhDC*5451::Tn10dTc[del-25] Δ*hin*-5717::FRT Δ*flhE*23076::FRT Δ*flgHI*23618::FKF | This study |
| EM15573 | *fliC*23165-*mneongreen* P$_{tetA}$*flhDC*5451::Tn10dTc[del-25] Δ*hin*-5717::FRT Δ*flgHI*23618::FRT / pTrc99AFF4 | This study |
| EM15574 | *fliC*23165-*mneongreen* P$_{tetA}$*flhDC*5451::Tn10dTc[del-25] Δ*hin*-5717::FRT Δ*flhE*23076::FRT Δ*flgHI*23618::FRT / pTrc99AFF4 | This study |
| EM15638 | *fliC*23165-*mneongreen* P$_{tetA}$*flhDC*5451::Tn10dTc[del-25] Δ*hin*-5717::FRT Δ*flgHI*23618::FRT Δ*flgE*22964::FRT / pTrc99AFF4 | This study |
| EM15639 | *fliC*23165-*mneongreen* P$_{tetA}$*flhDC*5451::Tn10dTc[del-25] Δ*hin*-5717::FRT Δ*flhE*23076::FRT Δ*flgHI*23618::FRT Δ*flgE*22964::FRT / pTrc99AFF4-*flgE*$^{+}$ | This study |
| EM14432 | Δ*araBAD*1001::*flgH* Δ*flgH*23616::FKF | This study |
| EM14433 | Δ*araBAD*1002::*flgI* Δ*flgI*23617::FKF | This study |
| EM14434 | Δ*araBAD*941::*flgHI* Δ*flgHI*23618::FKF | This study |
| EM14463 | Δ*flgH*7662 Δ*araBAD*1001::*flgH* | This study |
| EM14464 | Δ*flgI*7663 Δ*araBAD*1002::*flgI* | This study |
| EM14465 | Δ*flgHI*958 Δ*araBAD*941::*flgHI* | This study |
| EM808 | Δ*araBAD*1005::FRT | Lab collection |
| EM14678 | Δ*flgHI*958 P$_{tetA}$*flhDC*5451::Tn10dTc[del-25] / pTrc99AFF4 | This study |
| EM14676 | Δ*flgHI*958 P$_{tetA}$*flhDC*5451::Tn10dTc[del-25] Δ*flhE*23076::FCF / pTrc99AFF4 | This study |
| EM14750 | Δ*flgHI*958 P$_{tetA}$*flhDC*5451::Tn10dTc[del-25] Δ*flhE*23076::FCF Δ*flgE*23643::FKF / pTrc99AFF4 | This study |
| EM14751 | Δ*flgHI*958 P$_{tetA}$*flhDC*5451::Tn10dTc[del-25] Δ*flhE*23076::FCF Δ*flgE*23643::FKF / pTrc99AFF4-*flgE*$^{+}$ | This study |
| EM14752 | Δ*flgHI*958 P$_{tetA}$*flhDC*5451::Tn10dTc[del-25] Δ*flhE*23076::FCF Δ*flgJ*23644::FKF / pTrc99AFF4 | This study |
| EM14753 | Δ*flgHI*958 P$_{tetA}$*flhDC*5451::Tn10dTc[del-25] Δ*flhE*23076::FCF Δ*flgJ*23644::FKF / pTrc99AFF4-*flgJ*$^{+}$ | This study |
| EM14754 | Δ*flgHI*958 P$_{tetA}$*flhDC*5451::Tn10dTc[del-25] Δ*flhE*23076::FCF Δ*flgG*23645::FKF / pTrc99AFF4 | This study |
| EM14755 | Δ*flgHI*958 P$_{tetA}$*flhDC*5451::Tn10dTc[del-25] Δ*flhE*23076::FCF Δ*flgG*23645::FKF / pTrc99AFF4-*flgG*$^{+}$ | This study |
| EM14756 | Δ*flgHI*958 P$_{tetA}$*flhDC*5451::Tn10dTc[del-25] Δ*flhE*23076::FCF Δ*fliS*5720::FKF / pTrc99AFF4 | |
| EM14757 | Δ*flgHI*958 P$_{tetA}$*flhDC*5451::Tn10dTc[del-25] Δ*flhE*23076::FCF Δ*fliS*5720::FKF / pTrc99AFF4-*fliS*$^{+}$ | This study |
| EM14758 | Δ*fliC*7861::FRT Δ*hin*-5717::FCF (fliC$^{ON}$) P$_{tetA}$*flhDC*5451::Tn10dTc[del-25] Δ*flgH*23616::FKF / pTrc99AFF4 | This study |
| EM14759 | Δ*fliC*7861::FRT Δ*hin*-5717::FCF (fliC$^{ON}$) P$_{tetA}$*flhDC*5451::Tn10dTc[del-25] Δ*flgH*23616::FKF / pTrc99A FF4-*fliC*$^{+}$ | This study |
| EM14760 | Δ*fliC*7861::FRT Δ*hin*-5717::FCF (fliC$^{ON}$) P$_{tetA}$*flhDC*5451::Tn10dTc[del-25] Δ*flgH*23616::FKF / pTrc99A FF4-*fliC* (Δ aa 451-495) | This study |
| EM14761 | Δ*flhE*7404 Δ*fliC*7861::FRT Δ*hin*-5717::FCF (fliC$^{ON}$) P$_{tetA}$*flhDC*5451::Tn10dTc[del-25] Δ*flgH*23616::FKF / pTrc99AFF4 | This study |
| EM14762 | Δ*flhE*7404 Δ*fliC*7861::FRT Δ*hin*-5717::FCF (fliC$^{ON}$) | This study |

| | | |
|---|---|---|
| | $P_{tetA}$*flhDC*5451::Tn10dTc[del-25] Δ*flgH*23616::FKF / pTrc99A FF4-*fliC*$^{+}$ | |
| EM14763 | Δ*flhE*7404 Δ*fliC*7861::FRT Δ*hin*-5717::FCF (fliC$^{ON}$) $P_{tetA}$*flhDC*5451::Tn10dTc[del-25] Δ*flgH*23616::FKF / pTrc99A FF4-*fliC* (Δ aa 451-495) | This study |
| EM15551 | *fliC*23165-*mneongreen* $P_{tetA}$*flhDC*5451::Tn10dTc[del-25] Δ*hin*-5717::FRT Δ*flhE*23076::FRT Δ*flgBC*6557 | This study |
| EM15552 | *fliC*23165-*mneongreen* $P_{tetA}$*flhDC*5451::Tn10dTc[del-25] Δ*hin*-5717::FRT Δ*flgBC*6557 | This study |
| EM15553 | *fliC*23165-*mneongreen* $P_{tetA}$*flhDC*5451::Tn10dTc[del-25] Δ*hin*-5717::FRT Δ*flhE*23076::FRT Δ*flgJ*7600 | This study |
| EM15554 | *fliC*23165-*mneongreen* $P_{tetA}$*flhDC*5451::Tn10dTc[del-25] Δ*hin*-5717::FRT Δ*flgJ*7600 | This study |
| EM15555 | *fliC*23165-*mneongreen* $P_{tetA}$*flhDC*5451::Tn10dTc[del-25] Δ*hin*-5717::FRT Δ*flhE*23076::FRT Δ*flgG*7661 | This study |
| EM15556 | *fliC*23165-*mneongreen* $P_{tetA}$*flhDC*5451::Tn10dTc[del-25] Δ*hin*-5717::FRT Δ*flgG*7661 | This study |
| EM14439 | Δ*hin*-5717::FRT *fliC*5600$_{T237C}$ $P_{tetA}$*flhDC*5451::Tn10dTc[del-25] Δ*flgHI*23618::FKF | This study |
| EM14440 | Δ*hin*-5717::FRT *fliC*5600$_{T237C}$ Δ*flhE*7404 $P_{tetA}$*flhDC*5451::Tn10dTc[del-25] Δ*flgHI*23618::FKF | This study |
| EM4872 | Δ*hin*-5717::FRT *flgE*6506$_{S171C}$ $P_{tetA}$*flhDC*5451::Tn10dTc[del-25] | Lab collection |
| EM14769 | Δ*hin*-5717::FRT *flgE*6506$_{S171C}$ $P_{tetA}$*flhDC*5451::Tn10dTc[del-25] Δ*flhE*23076::FRT Δ*flgHI*23618::FKF | This study |
| EM2046 | Δ*hin*-5717::FRT *fliC*5600$_{T237C}$ $P_{tetA}$*flhDC*5451::Tn10dTc[del-25] | Lab collection |
| EM9889 | Δ*hin*-5717::FRT *fliC*23284$_{T237C, G426A}$ $P_{tetA}$*flhDC*5451::Tn10dTc[del-25] | This study |
| EM9890 | Δ*hin*-5717::FRT *fliC*23285$_{T237C, A427V}$ $P_{tetA}$*flhDC*5451::Tn10dTc[del-25] | This study |
| EM9891 | Δ*hin*-5717::FRT *fliC*23286$_{T237C, A449V}$ $P_{tetA}$*flhDC*5451::Tn10dTc[del-25] | This study |
| EM9892 | Δ*hin*-5717::FRT *fliC*23287$_{T237C, G426A, A449T}$ $P_{tetA}$*flhDC*5451::Tn10dTc[del-25] | This study |
| EM9893 | Δ*hin*-5717::FRT *fliC*23288$_{T237C, Q472L}$ $P_{tetA}$*flhDC*5451::Tn10dTc[del-25] | This study |
| EM15686 | Δ*hin*-5717::FRT *fliC*5600$_{T237C}$ $P_{tetA}$*flhDC*5451::Tn10dTc[del-25] Δ*flgHI*23618::FKF Δ*flhE*23076::FCF | This study |
| EM15706 | Δ*hin*-5717::FRT *fliC*23284$_{T237C, G426A}$ $P_{tetA}$*flhDC*5451::Tn10dTc[del-25] Δ*flgHI*23618::FKF Δ*flhE*23076::FCF | This study |
| EM15707 | Δ*hin*-5717::FRT *fliC*23285$_{T237C, A427V}$ $P_{tetA}$*flhDC*5451::Tn10dTc[del-25] Δ*flgHI*23618::FKF Δ*flhE*23076::FCF | This study |
| EM15708 | Δ*hin*-5717::FRT *fliC*23286$_{T237C, A449V}$ $P_{tetA}$*flhDC*5451::Tn10dTc[del-25] Δ*flgHI*23618::FKF Δ*flhE*23076::FCF | This study |
| EM15709 | Δ*hin*-5717::FRT *fliC*23287$_{T237C, G426A, A449T}$ $P_{tetA}$*flhDC*5451::Tn10dTc[del-25] Δ*flgHI*23618::FKF Δ*flhE*23076::FCF | This study |
| EM15710 | Δ*hin*-5717::FRT *fliC*23288$_{T237C, Q472L}$ $P_{tetA}$*flhDC*5451::Tn10dTc[del-25] Δ*flgHI*23618::FKF Δ*flhE*23076::FCF | This study |
| EM15447 | Δ*flgHI*958 $P_{tetA}$*flhDC*5451::Tn10dTc[del-25] *mneongreen-zapA* (N-ter, no linker) | This study |
| EM15505 | Δ*flgHI*958 $P_{tetA}$*flhDC*5451::Tn10dTc[del-25] *mneongreen-zapA* | This study |
| EM15918 | Δ*flgHI*958 $P_{tetA}$*flhDC*5451::Tn10dTc[del-25] *mneongreen-minD* (N-ter, SAGASA linker) | This study |
| EM15974 | Δ*flgHI*958 $P_{tetA}$*flhDC*5451::Tn10dTc[del-25] *mneongreen-minD* | This study |

**Table S2**: Strains used for FliK-Bla assay

| Name | Genotype | Reference |
|---|---|---|
| EM11563 | Δ*flhE*23076::FCF | This study |
| EM12322 | Δ*flhE*23076::FRT | Lab collection |
| TH13359 | *fliK7582::bla* (before stop) | Lab collection |
| TH13807 | *fliK7582::bla* Δ*flgA7656*(D6-214(of 219)) | Lab collection |
| TH13808 | *fliK7582::bla* Δ*flgB7657*(D6-133(of 138)) | Lab collection |
| TH13809 | *fliK7582::bla* Δ*flgC7658*(D6-129(of 134)) | Lab collection |
| TH13810 | *fliK7582::bla* Δ*flgD6540*(D2-220(of 232)) | Lab collection |
| TH13811 | *fliK7582::bla* Δ*flgE7659*(D6-398(of 403)) | Lab collection |
| TH13812 | *fliK7582::bla* Δ*flgF7660*(D6-246(of 251)) | Lab collection |
| TH13813 | *fliK7582::bla* Δ*flgG7661*(D6-255(of 260)) | Lab collection |
| TH13814 | *fliK7582::bla* Δ*flgH7662*(D6-227(of 232)) | Lab collection |
| TH13815 | *fliK7582::bla* Δ*flgI7663*(D6-360(of 365)) | Lab collection |
| TH13816 | *fliK7582::bla* Δ*flgJ7664*(D6-311(of 316)) | Lab collection |
| TH13817 | *fliK7582::bla* Δ*flgK7665*(D6-548(of 553)) | Lab collection |
| TH13818 | *fliK7582::bla* Δ*flgL7666(D6-312(of 317))* | Lab collection |
| TH27867 | *fliK7582::bla* Δ*flhE7404* | This study |
| TH27868 | *fliK7582::bla* Δ*flgA7656* Δ*flhE7404* | This study |
| TH27869 | *fliK7582::bla* Δ*flgB7657* Δ*flhE7404* | This study |
| TH27870 | *fliK7582::bla* Δ*flgC7658* Δ*flhE7404* | This study |
| TH27871 | *fliK7582::bla* Δ*flgD6540* Δ*flhE7404* | This study |
| TH27872 | *fliK7582::bla* Δ*flgE7659* Δ*flhE7404* | This study |
| TH27873 | *fliK7582::bla* Δ*flhE7404* | This study |
| TH27874 | *fliK7582::bla* Δ*flgG7661* Δ*flhE7404* | This study |
| TH27875 | *fliK7582::bla* Δ*flgH7662* Δ*flhE7404* | This study |
| TH27876 | *fliK7582::bla* Δ*flgI7663* Δ*flhE7404* | This study |
| TH27877 | *fliK7582::bla* Δ*flgJ7664* Δ*flhE7404* | This study |
| TH27878 | *fliK7582::bla* Δ*flgK7665* Δ*flhE7404* | This study |
| TH27879 | *fliK7582::bla* Δ*flgL7666* Δ*flhE7404* | This study |

#### **Table S3**: Plasmids used in this study

| Name | Genotype and usage | Source or Reference |
|---|---|---|
| pKD46 | Lambda Red | 2 |
| pWRG730 | Lambda Red | 3 |
| pCP20 | FRT flipping | 4 |
| pKD4 | FKF cassette insertion | 2 |
| pEM8731 | pKH70-$P_{rpsM}$-*mneongreen*<br>Visualisation cells. Constitutive promoter. | 5 |
| pEM8313 | pKH70-$P_{rpsM}$-*mcerulean*<br>Visualisation cells. Constitutive promoter | 5 |
| pEM8913 | pKH70-$P_{rpsM}$-*flhE*<br>Complementation *flhE.* Constitutive promoter | This study |
| pXY027 | pCA24N-*ftsZ-gfp* (C-terminal, non-functional) | Addgene Plasmid #98915<br>1 |
| pEM12582 | pCA24N- *mreB-mneongreen-mreB* (Between aa G228-D229, SAGASA linker)<br>Visualisation of MreB by confocal microscopy | This study |
| pTrc99AFF4 | Empty vector, IPTG inducible promoter | 6 |
| pTrc99A FF4-*fliC*$^{+}$ | Expression of *fliC* full length (CDS aa 1 – aa 495) | Lab collection |
| pEM14674 /<br>pTrc99A FF4-*fliC*<br>(∆ aa 451-495) | Expression of *fliC* C-terminal truncated (CDS aa 1 – aa 450) | This study |
| pTrc99AFF4-*flgE*$^{+}$ | Expression of hook subunits *flgE* | Lab collection |
| pTrc99AFF4-*flgJ*$^{+}$ | Expression of rod cap *flgJ* | Lab collection |
| pTrc99AFF4-*flgG*$^{+}$ | Expression of distal rod subunit *flgG* | Lab collection |
| pTrc99AFF4-*fliS*$^{+}$ | Expression of flagellin chaperone *fliS* | Lab collection |
| pEM12581 | pCA24N- *mneongreen-minD* (N-terminal, SAGASA linker)<br>Construction of MinD-mNeonGreen | This study |

### **Table S4**: Primers used in this study

| Number | Name | Sequence (5' -> 3') | Usage |
|---|---|---|---|
| 3011 | Δ*flhE*-FCF-fw | gaccattggaggaaaataatgcgtaaatggctggcgttGTGTAGGCTGGAGCTGCTTC | Deletion |
| 3012 | Δ*flhE*-FCF-rv | gggcggttagcggtagttcacaatcacctgattgctccgCATATGAATATCCTCCTTAG | Deletion |
| 3028 | 5'-*flhE*-seq-fw | GGTATTACTGGTTAACCATGCG | Sequencing |
| 3029 | 3'-*flhE*-seq-rv | ACAGTAAAAGCTAAGCTGGG | Sequencing |
| 3876 | 5'-*flhE*-Cter-fw | ATGCGTAAATGGCTGGCGTT | Sequencing |
| 3877 | 3'-*flhE*-Nter-rev | TTAGCGGTAGTTCACAATCACC | Sequencing |
| 3407 | 5'-EcoRI-*flhE*-fw | ggacagaattcataaggaggaaaaacatATGCGTAAATGGCTGGCGTT | Cloning |
| 3408 | 3'-NotI-*flhE*-rev | gtactgcggccgcTGCAGTTTTACCGCCGACGC | Cloning |
| 4052 | 5'-*KanSceI-flhE*-del-Nter-fw | gtcatatccgcatgacggcgaccattggaggaaaataatgAGGGTTTTCCCAGTCACGAC | Deletion |
| 4053 | 3'-*KanSceI-flhE*-del-Nter-rev | ttacacccataccgctatcctgccacgcgccttcgcctgcTGCTTCCGGCTCGTATGTTG | Deletion |
| 4684 | 5'-flhBA-FKF-fw | gtggcagaagagagcgacgacgacaaaacagaagcccccaGTGTAGGCTGGAGCTGCTTC | Deletion |
| 4685 | 3'-flhB-FKF-rev | tccagcgcttctggcaccggaaggttctcaggttgtggagCATATGAATATCCTCCTTAG | Deletion |
| 4686 | 5'-flhA-FKF-fw | ggtcgcgatgctgcgcctgcccagcaacctgaaatcgacgGTGTAGGCTGGAGCTGCTTC | Deletion |
| 4687 | 3'-flhBA-FKF-rev | atatgacggttatcggaaagctcaaggttcgacaacaccaCATATGAATATCCTCCTTAG | Deletion |
| 4688 | 5'-DfliPQR-FRT-fw | ggagatcctgatgcgccgtttgttattcctttctctggcgGTGTAGGCTGGAGCTGCTTC | Deletion |
| 4689 | 3'-DfliP_aa10-241-FRT-rev | gattcaggagtcattttgcgcctctaactgtaaaagctttCATATGAATATCCTCCTTAG | Deletion |
| 4690 | 3'-DfliPQR-FRT-rev | ttatgggttattatttatcggcatctcgctaacaatatcaCATATGAATATCCTCCTTAG | Deletion |
| 5400 | 5'-G228-mreB-overlapping | ATGTTGAAAAAATTTCGTGG | MreB-mNG Gibson Assembly |
| 5401 | 3'-G228-mreB-overlapping | tgctgatgcgcctgcacttgaGCCCGGATAAGCGGAACCGA | MreB-mNG Gibson Assembly |
| 5402 | 5'-mNG-overlapping | tcaagtgcaggcgcatcagcaATGGTATCGAAGGGCGAGGA | MreB-mNG Gibson Assembly |
| 5404 | 3'-mNG-Venus-overlapping | ggccgaagcacctgcggaaccTTTATACAGTTCATCCATGC | MreB-mNG Gibson Assembly |
| 5405 | 5'-D229-*mreB*-overlapping | ggttccgcaggtgcttcggccGACGAAGTCCGCGAGATCGA | MreB-mNG Gibson Assembly |
| 5406 | 3'-D229-*mreB*-overlapping | CTACTCTTCGCTGAACAGGT | MreB-mNG Gibson Assembly |
| 5407 | 5'-pXY027-*mreB*-fw-GA | acaggtcattggaaaacatgccacgaaatttttcaacatACTAGTAGTTAATTTCTCCT | MreB-mNG Gibson Assembly |
| 5408 | 3'-pXY027-*mreB*-rev-GA | atgatcgatatgcacggcggcgacctgttcagcgaagagtagGCGGCCGCTAAGGGTCGA | MreB-mNG Gibson Assembly |
| 5409 | 5'-mNG-*minD*-GA-fw | tttcacacagaattcattaaagaggagaaattaactactagtATGGTATCGAAGGGCGAG | MinD-mNG Gibson Assembly |
| 5410 | 3'-mNG-*minD*-GA-rev | acaacaataatgcgtgccattgctgatgcgcctgcactTTTATACAGTTCATCCAT | MinD-mNG Gibson Assembly |
| 5411 | 5'-*minD*-GA-fw | agtgcaggcgcatcagcaATGGCACGCATTATTGTTGT | MinD-mNG Gibson Assembly |
| 5412 | 3'-*minD*-GA-rev | cagctaattaagcttggctgcaggtcgacccttagcggccgcTTATCCTCCGAACAGGCG | MinD-mNG Gibson Assembly |
| 5413 | 5'-pXY027-*minD*-GA-fw | gcggccgctaagggtcgacc | MinD-mNG Gibson Assembly |
| 5414 | 3'-pXY027-*minD*-GA-rev | actagtagttaatttctcct | MinD-mNG Gibson Assembly |
| 2522 | *fliC*-G426-*tetRA*-FW | gctgctttggcacaggttgacacgttacgttctgacctggTTAAGACCCACTTTCACATT | *fliC* point mutant |
| 2523 | *fliC*-A449-*tetRA*-RV | aaacttcggtcgcgtagtcggaatcttcgatacggctacgCTAAGCACTTGTCTCCTG | *fliC* point mutant |
| 2524 | *fliC*-G426A-FW | gctgctttggcacaggttgacacgttacgttctgacctggCtgcggtacagaaccgtttc | *fliC* point mutant |
| 3751 | 5'-*fliC*-A427V-fw | gctgctttggcacaggttgacacgttacgttctgacctgggtgTggtacagaaccgtttc | *fliC* point mutant |
| 2526 | *fliC*-1484-RV | cgcagtaaagagaggacgtttt | *fliC* point mutant |
| 3752 | 3'-*fliC*-A449T-rev | aacttcggtcgcgtagtcggaatcttcgatacggctacgCGTagaagtcaggttgtttac | *fliC* point mutant |
| 2525 | *fliC*-A449V-RV | aacttcggtcgcgtagtcggaatcttcgatacggctacggAcagaagtcaggttgtttac | *fliC* point mutant |
| 3753 | 3'-*fliC*-Q472L-rev | cctggttcgcctgcgccagaacggaggtaccggcctgcagCAGAATCTGCGCGCGAGACA | *fliC* point mutant |
| 2527 | *fliC*-1147-FW | gcaagtaaagccgaaggtca | *fliC* point mutant |
| 202 | 5'-fliCend-tetR_fw | ggcgaaccaggttccgcaaaacgtcctctctttactgcgtTTAAGACCCACTTTCACATT | *fliC* transcriptional fusion |
| 203 | 3'-fliCend-3'UTR-tetA_rv | tgccttgattgtgtaccacgtgtcggtgaatcaatcgccggaCTAAGCACTTGTCTCCTG | *fliC* transcriptional fusion |
| 3395 | 5'-mNeonGreen-*fliC*-trans | gaaccaggttccgcaaaacgtcctctctttactgcgttaaGAATTCATAAGGAGGAAAAA | *fliC* transcriptional fusion |
| 3396 | 3' -mNeonGreen-*fliC*-trans | ccttgattgtgtaccacgtgtcggtgaatcaatcgccggaTTATCATTTATACAGTTCAT | *fliC* transcriptional fusion |
| 619 | *flhDC*-P5x_fw_52 | CATAAGACACTTTTTACACACCAC | $P_{pro/syn}$-*flhDC* |
| 3718 | *3'-PproC-flhD_rv* | tgtgtagagggaaaccgttgtggtctccctgaatatATCATAcgagccttatgcatgccc | $P_{pro/syn}$-*flhDC* |
| 3719 | 3'-PproB-flhD_rv | tgtgtagagggaaaccgttgtggtctccctgaatatATATTAcgagccttatgcatgccc | $P_{pro/syn}$-*flhDC* |
| 3720 | *3'-PproA-flhD_rv* | tgtgtagagggaaaccgttgtggtctccctgaatatAGCCTAcgagccttatgcatgccc | $P_{pro/syn}$-*flhDC* |
| 3721 | 3'-Ppro1-flhD_rv | tgtgtagagggaaaccgttgtggtctccctgaatatAGATACcgagccttatgcatgccc | $P_{pro/syn}$-*flhDC* |
| 3251 | *5'dPflhDC(-598)-proD_ fw* | actttttacacaccacccggatgcttcatttaaatgggtgCAGCTAACACCACGTCGTCC | $P_{pro/syn}$-*flhDC* |
| 3255 | 3'-RBS(proD)-flhDC_rv | ATAAATGTGTTTTAGCAACTCGGATGTATGCATTGTTCCCATctagtaCTTTCCTGTGTG | $P_{pro/syn}$-*flhDC* |
| 2884 | FliK*bla*-fw | caggccgccgcgcgcggcaatggcgcagtggatatcttcgcccacccagaaacgctggtg | FliK-Bla fusion |
| 2885 | FliK*bla*-Rev | gcgtggaaaagacgcggataatcatgctacctctggcgttaccaatgcttaatcagtgag | FliK-Bla fusion |

#### Supplementary References


1. Buss, J. *et al.* A multi-layered protein network stabilizes the *Escherichia coli* FtsZ-ring and modulates constriction dynamics. *PLoS Genet* **11**, e1005128 (2015).
2. Datsenko, K. A. & Wanner, B. L. One-step inactivation of chromosomal genes in *Escherichia coli* K-12 using PCR products. *Proceedings of the National Academy of Sciences* **97**, 6640–6645 (2000).
3. Blank, K., Hensel, M. & Gerlach, R. G. Rapid and Highly Efficient Method for Scarless Mutagenesis within the *Salmonella enterica* Chromosome. *PLOS ONE* **6**, e15763 (2011).
4. Cherepanov, P. P. & Wackernagel, W. Gene disruption in *Escherichia coli* : TcR and KmR cassettes with the option of Flp-catalyzed excision of the antibiotic-resistance determinant. *Gene* **158**, 9–14 (1995).
5. Delgadillo-Guevara, M., Halte, M., Erhardt, M. & Popp, P. F. Fluorescent tools for the standardized work in Gram-negative bacteria. 2024.01.18.576257 Preprint at https://doi.org/10.1101/2024.01.18.576257 (2024).
6. Ohnishi, K., Fan, F., Schoenhals, G. J., Kihara, M. & Macnab, R. M. The FliO, FliP, FliQ, and FliR proteins of *Salmonella* Typhimurium: putative components for flagellar assembly. *J Bacteriol* **179**, 6092–6099 (1997).
