## Supplementary material for "FlhE functions as a chaperone to prevent formation of periplasmic flagella in Gram-negative bacteria": Video S1 legend

Video S1: Phase contrast time-lapse imaging showing 3 exemplary cells (P*_tetA_*-*flhDC* ∆*flgHI* ∆*flhE)* after 2,5 hours induction of P*_tetA_*-*flhDC* with AnTc. Some cells are able to rotate their cell body, either attached to the coverslip (left panel), rotating in a circular motion (middle panel) or rotating the cell body in a similar manner to bacterial cell from the *Leptospira* genus. Scale bar = 10 μm and time in milliseconds are indicated (43 msec /frame).
